# Glycans on the SARS-CoV-2 Spike Control the Receptor Binding Domain Conformation

**DOI:** 10.1101/2020.06.26.173765

**Authors:** Rory Henderson, Robert J Edwards, Katayoun Mansouri, Katarzyna Janowska, Victoria Stalls, Megan Kopp, Barton F. Haynes, Priyamvada Acharya

## Abstract

The glycan shield of the beta-coronavirus (β-CoV) Spike (S) glycoprotein provides protection from host immune responses, acting as a steric block to potentially neutralizing antibody responses. The conformationally dynamic S-protein is the primary immunogenic target of vaccine design owing to its role in host-cell fusion, displaying multiple receptor binding domain (RBD) ‘up’ and ‘down’ state configurations. Here, we investigated the potential for RBD adjacent, N-terminal domain (NTD) glycans to influence the conformational equilibrium of these RBD states. Using a combination of antigenic screens and high-resolution cryo-EM structure determination, we show that an N-glycan deletion at position 234 results in a dramatically reduced population of the ‘up’ state RBD position. Conversely, glycan deletion at position N165 results in a discernable increase in ‘up’ state RBDs. This indicates the glycan shield acts not only as a passive hinderance to antibody meditated immunity but also as a conformational control element. Together, our results demonstrate this highly dynamic conformational machine is responsive to glycan modification with implications in viral escape and vaccine design.

## Introduction

The ongoing SARS-CoV-2 (SARS-2) pandemic presents an urgent need for the development of a protective vaccine. The primary immunogenic target for the vaccines in development is the viral transmembrane S-protein trimer. Each protomer of the trimer is split into an N-terminal receptor binding S1 subunit and a C-terminal fusion element containing S2 subunit. The S1 subunit is further split into an N-terminal domain (NTD), two subdomains (SD1 and SD2) as well as the receptor binding domain (RBD) that together cap the conserved elements of the S2 subunit. The fusion event is marked by the shedding of the S1 subunit and large conformational transitions in the S2 subunit. The necessity to maintain a large free energy gradient between the prefusion, immune protective state of the molecule and the post-fusion state results in a highly dynamic macromolecular structure. The S1 subunit is particularly dynamic, presenting the RBD in two distinct states: a receptor binding site occluded ‘down’ state in which the RBDs rest against their adjacent protomer’s NTD, and a receptor binding site exposed ‘up’ state. It is this RBD ‘up’ state to which the majority of neutralizing responses are observed in convalescent SARS-2 infected individuals^1^ As conformational evasion is a well-known virus escape mechanism, it is critical to understand the mechanism by which these dynamics are controlled.

Structural studies of the β-CoV S-protein have focused primarily on a soluble, ectodomain construct with and without stabilizing proline mutations (2P). This includes structures for SARS-2^1,2^, SARS^3–7^, MERS^3,8^, and other human^9,10^ and murine^11^ β-CoV ectodomains. Structures for the SARS and MERS ectodomains revealed the presence of one and two RBD ‘up’ states with a three RBD ‘up’ state observed in the MERS ectodomain demonstrating the breadth of RBD configurations available to the spike. Interestingly, these states were not observed in the human β-CoVs HKU1 and OC43 nor in a Murine β-CoV, suggesting mutations in the spike protein can confer dramatic differences in the propensity of the RBD to sample its available conformational space.

Our quantitative examination of the available β-CoV S-protein structures recently revealed the S1 and S2 subunit domains of different β-CoV viruses occupy a diverse array of configurations^12^. Based upon this analysis we predicted the S-protein conformation was particularly sensitive to mutations at the interfaces between domains and subunits. Indeed, mutations at these sites had major impacts on the configuration of the protein, especially on the RBD ‘up’/’down’ distribution^12^. While these and other studies ^13,14,15^ have demonstrated the role of protein-protein contacts in determining the conformation of the S-protein, the influence on RBD configuration of glycosylation at or near interfacial domain regions is poorly understood. Like other class I viral fusion proteins, the β-CoV S-proteins are heavily glycosylated, obscuring the spike surface and limiting the targetable area for immune responses. A recent site-specific analysis of the glycosylation patterns of the SARS-2 S-protein revealed extensive variation in the glycan type, indicating marked differences in processing enzyme accessibility at each site^16^. Together, the wide variation in spike conformation coupled with the presence of glycans adjacent to the RBD suggests, among the many factors affecting the RBD position, glycosylation patterns may provide a means by which to control its conformational equilibrium.

In this study we have investigated the potential for two SARS-2 NTD glycans in close proximity to the RBD to influence the conformational distribution of the RBD ‘up’ and ‘down’ states. Analysis of the available SARS-2 ‘up’ state structures suggested N165 and N234 glycans may interact with the ‘up’ state RBD acting as both direct stabilizers of the ‘up’ state and as steric blocks to transitions to the ‘down’ state. We combined binding studies by surface plasmon resonance, with structural studies using negative stain electron microscopy (NSEM) and single-particle cryo-electron microscopy (cryo-EM) to define shifts in the ‘up’/’down’ state equilibrium in glycan-deleted mutants of the SARS-2 spike ectodomain. Together, our results demonstrate that RBD proximal glycans can influence the propensity of the S-protein adopt multiple configurations suggesting a means for viral escape and therefore the need to consider non-RBD neutralizing responses in vaccine design.

## Results

### Structure analysis identifies glycans with the potential to modify the S-protein conformation

In order to establish whether glycans may indeed alter the RBD orientation, we first examined the SARS-2 glycan density at positions 165 and 234 in the cryo-EM maps from three previously published SARS-2 structures^2,17^. In the ‘down’ state, the N234 glycan resides in a cleft formed by the NTD and RBD (Supplemental Figure 1A) while in the ‘up’ state, it occupies the region of the RBD ‘down’ state (Figure 1A). This suggests that the solvated ‘up’ state configuration is preferred and must be shifted in order to accommodate the ‘down’ state. The presence of this glycan may act as a hinderance to ‘up’-to-’down’ state transitions while sterically hindering the ‘down’ state by limiting RBD to NTD packing. An additional glycan at N165 residing toward the apical position of the NTD is in close proximity to the RBD and therefore may also influence the RBD position. Unlike the N234 glycan, the position of the N165 glycan presents no apparent restriction to the RBD positioning in the ‘down’ state (Supplemental Figure 1B). However, clear density for this glycan is observed occupying the region the RBD rest in the closed state, potentially forming interactions with the ‘up’ state RBD (Figure 1A). This suggests this glycan may act to stabilize the RBD ‘up’ state. Alternatively, its presence near the RBD in the ‘down’ state may confer a degree of stability to the fully closed state. Together, these observations combined with our recent results suggesting remarkable conformational sensitivity to mutations suggest these glycans may act to stabilize the observed RBD ‘up’/’down’ equilibrium. We next asked whether RBD proximal NTD glycans occur in other β-CoVs for which high-resolution structural data is available. For this, we examined structures for MERS, SARS, OC43, HKU1, and a Murine β-CoV S-protein ectodomains (PDB IDs 5W9H, 6CRW, 6OHV, 5I08, and 3JCL, respectively), identifying three MERS (N155, N166, and N236), two SARS-2 synonymous SARS (N158 and N227), and one OC43 (N133), and two HKU1 (N132 and N19) glycosylation sites proximal to their respective RBDs (Figure 1B). No RBD adjacent glycosylation sites were observed in the Murine S-protein. While the MERS and SARS glycans display similar extensions into the RBD space in the one ‘up’ state, the lack of ‘up’ state structures for OC43 and HKU1 precludes examination of their RBD adjacent NTD glycans in the absence of the RBD. However, the HKU1 N132 and OC43 N133 glycans extend upward, away from the RBD suggesting these glycans are unlikely to exert a strong influence on the RBD conformation (Supplemental Figure 1C). Interestingly, while cryo-EM reconstructions for SARS-2, MERS, and SARS yield ‘up’ state RBDs, these states were not reported for any of the OC43, HKU1, or Murine datasets. Together, these observations suggest RBD proximal NTD glycans may indeed affect the conformational distribution of ‘up’/’down’ RBD states.

**Figure 1.**
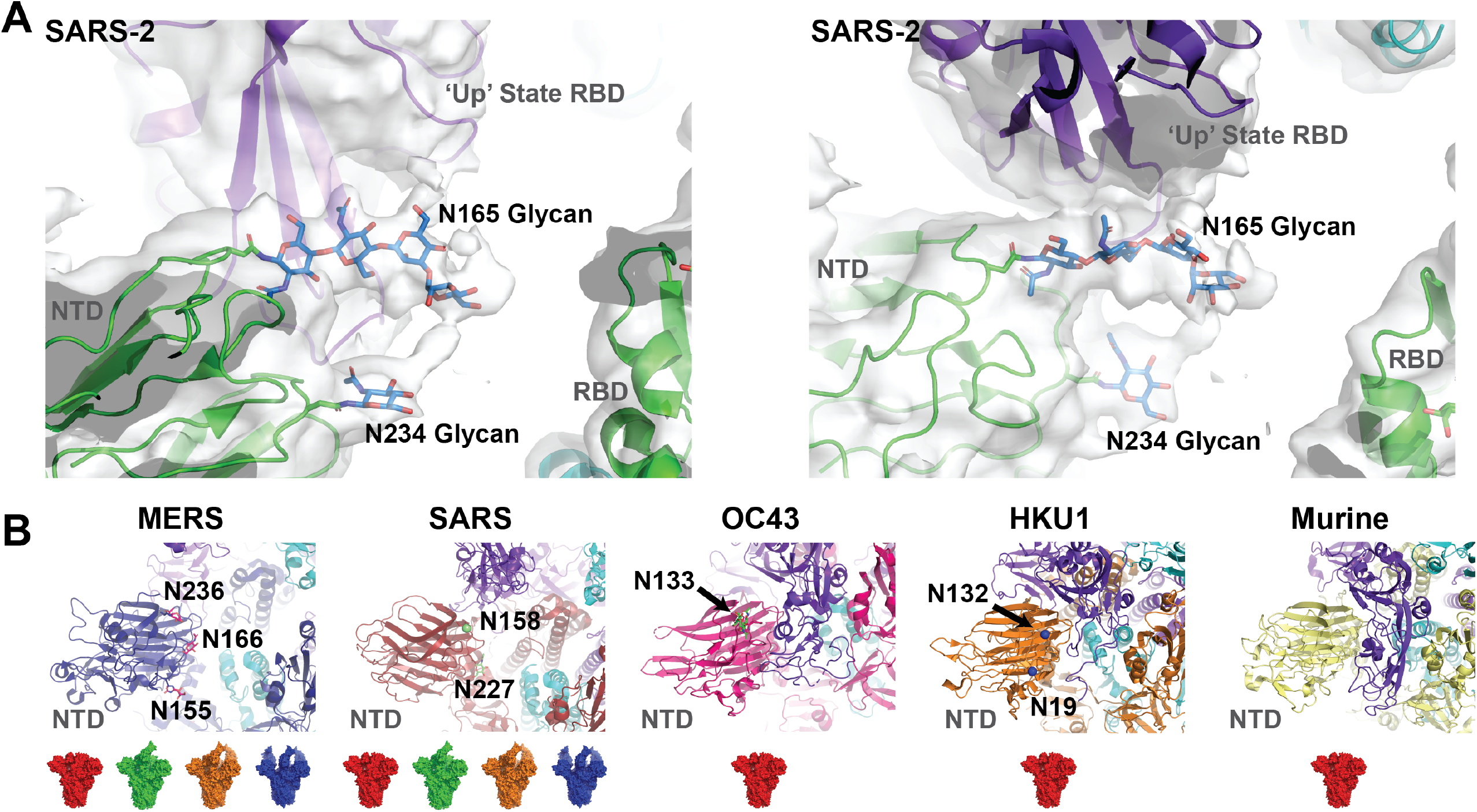
RBD proximal NTD glycans of SARS-2 MERS, SARS, and other β-CoV S-proteins. **A)** (*left*) Side view of the one RBD ‘up’ state SARS-2 structure and map (PDB ID 6VSB; EMDB 21375) depicting the reconstructed N165 and N234 NTD glycans protruding into the space occupied by the RBD in the ‘down’ state. (right) top view of the N165 and N234 glycans. **B)** Structures of the MERS, SARS, OC43, HKU1, and Murine S-proteins (PDB IDs 5W9H, 6CRW, 6OHV, 5I08, and 3JCL, respectively) depicting the location of RBD proximal N-linked glycans. Closed (red), 1-up (green), 2-up (orange), and 3-up (blue) RBD state surfaces below cartoon representations indicate whether such states have been observed for each timer.

### RBD conformation and antigenicity of the N-glycan deleted S-proteins reveals differential stabilization of RBD ‘up’ and ‘down’ states

In order to examine the extent to which the N234 and N165 glycans influence the conformational distribution of the S-protein, we produced di-proline (2P) stabilized^8^ S-protein ectodomain^2^ N234A and N165A mutants. The protein yields after StrepTactin purification were 2.0 mg and 0.8 mg per 1L culture supernatant, respectively for the N234A and the N165A mutant. (Supplemental Figures 2 and 3). To assess the reactivity of the glycan-deleted spike ectodomain mutants to the ACE-2 receptor, we tested binding of the spike to an ACE-2 ectodomain construct bearing a C-terminal mouse Fc tag immobilized on an anti-Fc surface. SPR binding assays showed that while the N165A mutant displayed ~10-20% increased binding levels to the unmutated construct, the N234A mutant showed a decrease of ~50-60% (Figure 2A and Supplemental Figure 2B). Because ACE-2 binding requires the RBD be in the up position, the SPR data suggests that the N165A mutant displays more ‘up’ (or open) states, whereas the N2345A mutant displays more ‘down’ (or closed) states.

**Figure 2.**
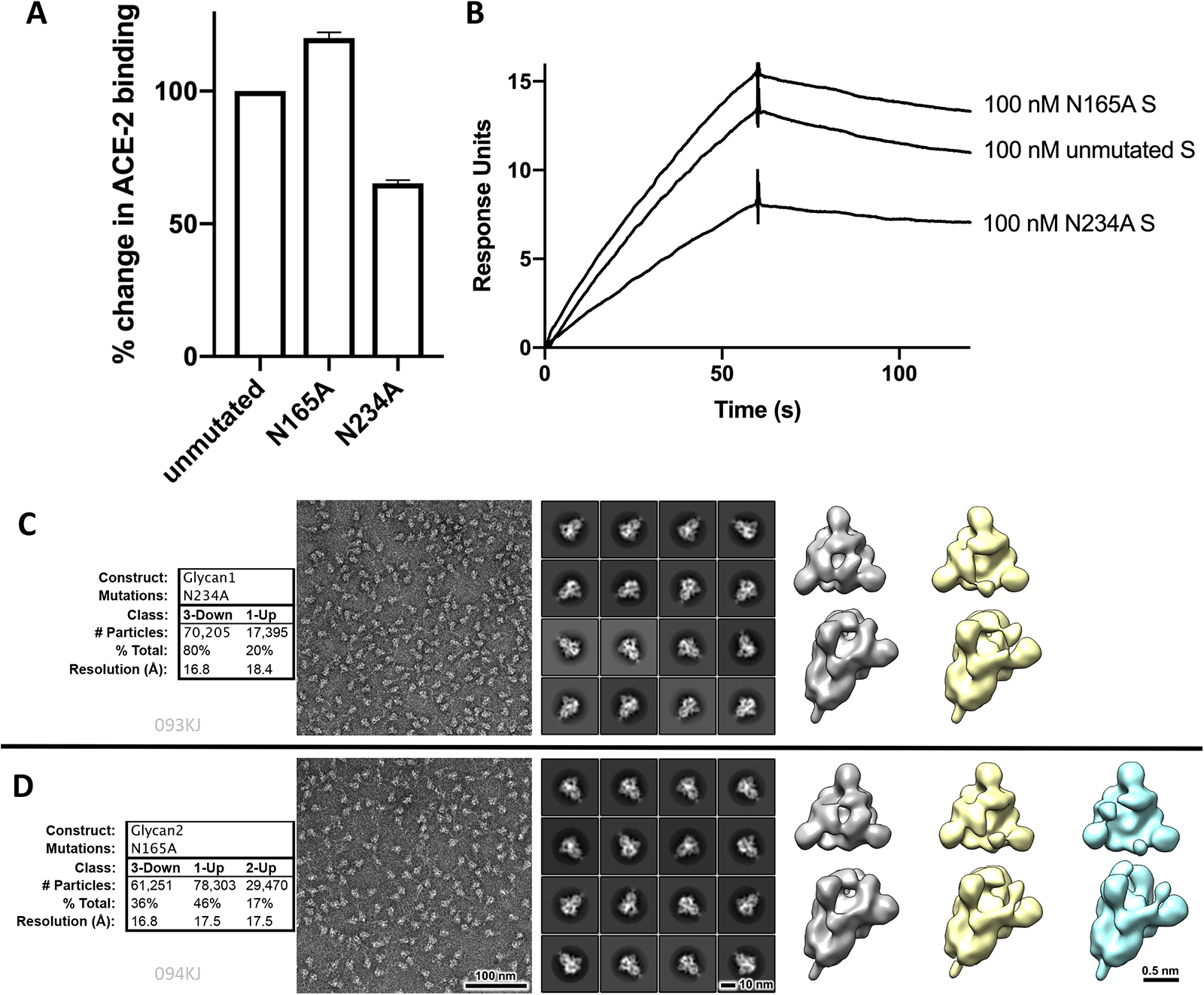
Structure and antigenicity of the N165A and N234A SARS-CoV-2 ectodomain spikes. **A)** Percentage change in ACE2 binding for the N234A, and N165A mutant spikes, relative to the unmutated spike. Binding was measured by SPR with ACE-2 (with a C-terminal Fc tag) captured on on an anti-Fc surface, and the spike as analyte. Error bars represent results from four independent injections. **B)** Representative ACE2 binding SPR response curves. **C)** NSEM results for the N234A spike. **D)** NSEM results for the N165A spike. For figures (C) and (D) shown from left to right are percentages of discrete 3D populations observed, representative micrograph, representative 2D class averages, discrete populations obtained by 3D classification.

We next examined the ‘up’/’down’ state distribution of both mutants via negative stain electron microscopy (NSEM). Heterogenous classification of the N234A mutant particles revealed a dramatic shift from a ~1:1 ‘up’ v. ‘down’ state distribution in the unmutated 2P^12,17^ to a ratio of ~1:4 in the down state (Figure 2A). Remarkably, the N165A mutant shifted the distribution in the opposite direction, displaying a higher propensity to adopt RBD “up” states yielding a ~2:1 - ‘up’ state to ‘down’ state ratio, with ~ 17% of the “up’ population being a 2-RBD “up” class (Figure 2A). Together, the ACE-2 binding and the NSEM results demonstrated that both NTD N-glycan deletions have distinct impacts on the RBD distribution.

### High-resolution cryo-EM structures of the N-glycan deleted constructs indicates modest perturbation to S-protein configuration

We next turned to cryo-EM for high resolution structure determination to visualize the impact of the glycan deletions on the local and global configuration of the S-protein domains. We collected and processed 7,269 and 8,068 images for the N165A and N234A mutant, respectively, to yield particle stacks cleaned by 2D classification and multiple rounds of *ab initio* classification and heterogenous refinement in cryoSPARC^18^ using 20 Å low pass filtered ‘up’ state and ‘down’ state maps generated from available SARS-2 structures. Initial maps for high resolution refinement were generated from sorted particles via *ab initio* reconstruction (Supplemental Figures 4 and 5). The resulting particle distribution for the N234A mutant was predominantly ‘down’ with a minor, ~6%, ‘up’ state population while that of the N165A mutant was ~50% ‘down’ and 50% one ‘up’ as was observed for the unmutated spike previously^12,17^. We were unable to identify a particle subset corresponding to a two ‘up’ state in the cryo-EM dataset. The ‘up’/’down’ state populations obtained via NSEM for unmutated^12^, glutaraldehyde fixed SARS-2 S-protein ectodomain match the previously observed cryo-EM distribution^17^. Here, using the same approach, we find that these distributions are dramatically and differentially shifted with mutation of either the N165 or N234 to alanine with the SPR, NSEM, and cryo-EM distribution tracking in the same direction with the exception of the N165A cryo-EM particles for which a two RBD ‘up’ state was not observed. Considering the concordance between the SPR and NSEM results, this may be due to particle processing and the potential for a relatively disordered ‘up’ state RBDs in the two ‘up’ state with the glycan deletion.

We next examined the high-resolution details of the cryo-EM maps. Refinement of the N234A mutant ‘down’ state using C3 symmetry resulted in a 3.0 Å map with coordinates fit to this map yielding a structure aligning to the unmutated 2P structure (PDB ID 6VXX) with a ~0.6 Å RMSD. Alignment of the S2 subunit revealed the structures to be nearly identical in these regions (RMSD ~0.4 Å). Examination of the NTD to RBD interface using this alignment revealed a shift of the NTD toward the RBD (Figure 3A-D). Weak density for the N165 glycan was observed suggestive of an overall similar position relative to that observed previously (Figure 3B and C) The one RBD ‘up’ state map was refined to 4.8 Å resolution using C1 symmetry. Comparison of the one RBD ‘up’ state structure fit to this map to its unmutated counterpart (PDB ID 6VYB) suggests a slight shift of the RBD with the N234A mutation (Figure 3E and F). However, the limited resolution of this structure limits close examination of this movement. Nevertheless, density for the N165 glycan was observed for the NTD adjacent to the vacant RBD site (‘up’ adjacent) and for the NTD glycan adjacent to the ‘down’ state RBD proximal to the vacant site (‘down’ free). Each occupies a configuration consistent with previous observations in the unmutated form. Interestingly, clear density for the N165 glycan is not observed for the NTD adjacent to the ‘down’ state RBD contacting the ‘up’ state RBD (‘down’ adjacent). Together, the structures show that the while clear differences between the unmutated and N234A mutant are observed, the overall configuration of the structures are similar to their respective unmutated counterparts. These differences do not appear to have significant impacts on the N165 glycan configuration.

**Figure 3.**
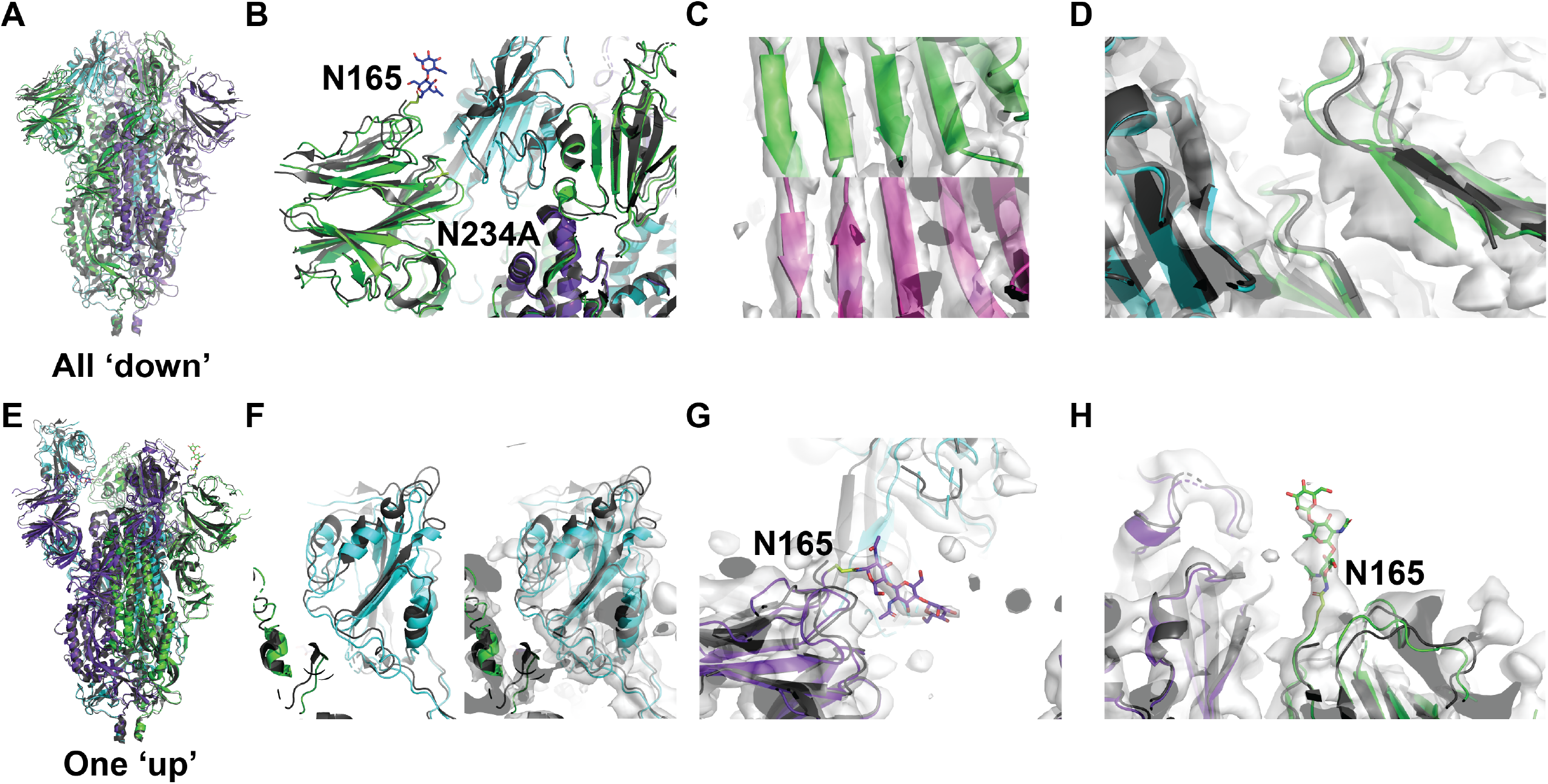
Structural comparison of the N234A mutant in the ‘up’ and ‘down’ configurations to the unmutated spike. **A)** Side view of the symmetric ‘down’ state N234A mutant S-protein trimer aligned to the unmutated trimer (PDB ID 6VXX, grey). **B)** Side view of a ‘down’ state NTD (green) depicting the shifted NTD relative to the unmutated form (grey) and the N165 glycan. Adjacent RBD is colored cyan. **C)** (*upper*) Map view of the apical β-sheet motif of the NTD for the N234A ‘down’ state (*lower*) Map view of the apical β-sheet motif of the NTD from the unmutated ‘down’ state (PDB ID 6VXX) **D)** The N234A trimer map density with the NTD (green) and RBD (cyan) coordinates aligned to the unmutated form (grey). **E)** Side view of the ‘up’ state N234A mutant S-protein trimer aligned to the unmutated trimer (PDB ID 6VYB, grey). **F)** (*left*) Cartoon representation of N234A ‘up’ state RBD (cyan) relative to the unmutated ‘up’ state RBD (grey). (*right*) as in (*left*) with the map density. **G)** Side view of the N165 glycan extending into the RBD ‘down’ state region near the ‘up’ state RBD. **H)** Top view of the N165 glycan extending into the RBD ‘down’ state region near the ‘up’ state RBD.

Refinement of the N165A ‘up’ and ‘down’ states resulted in maps with resolutions of 3.6 Å using C1 symmetry and 3.3 Å using C3 symmetry, respectively. Similar to the N234A mutant, the N165A mutant structures showed an overall similar arrangement of the various domains. Alignment of the ‘down’ state structure of the N165A mutant with that of the unmutated spike yielded an RMSD of 0.81 Å with an S2 subunit alignment RMSD of 0.36 Å. Unlike the N234A mutant, the N165A mutant NTD is shifted away from the adjacent RBD (Figure 4A-D). Interestingly, clear density for the N234 glycan was not observed. The one ‘up’ state structure of the N165A mutant displayed a similar (Supplemental Figure 6), albeit slightly less shifted, arrangement of the NTD in the ‘down’ adjacent protomer (Supplemental Figure 6B). This shift in not observed in either of the other two NTDs indicating the NTD shift is sensitive to S1 and S2 subunit arrangements (Supplemental Figure 6C and D). The 1-’up’ RBD resides in largely the same position as that of the unmutated spike with only minor differences due potentially to the lower relative resolution of this region (Figure 4E-H). Density for the N234 glycan was not observed for any of the protomers, consistent with the ‘down’ state map. Together, the results of the N165A and N234A structural analysis results suggest that these two glycans play a differential role in influencing the SARS-CoV-2 RBD arrangement, shifting the NTD toward or away from the adjacent RBDs.

**Figure 4.**
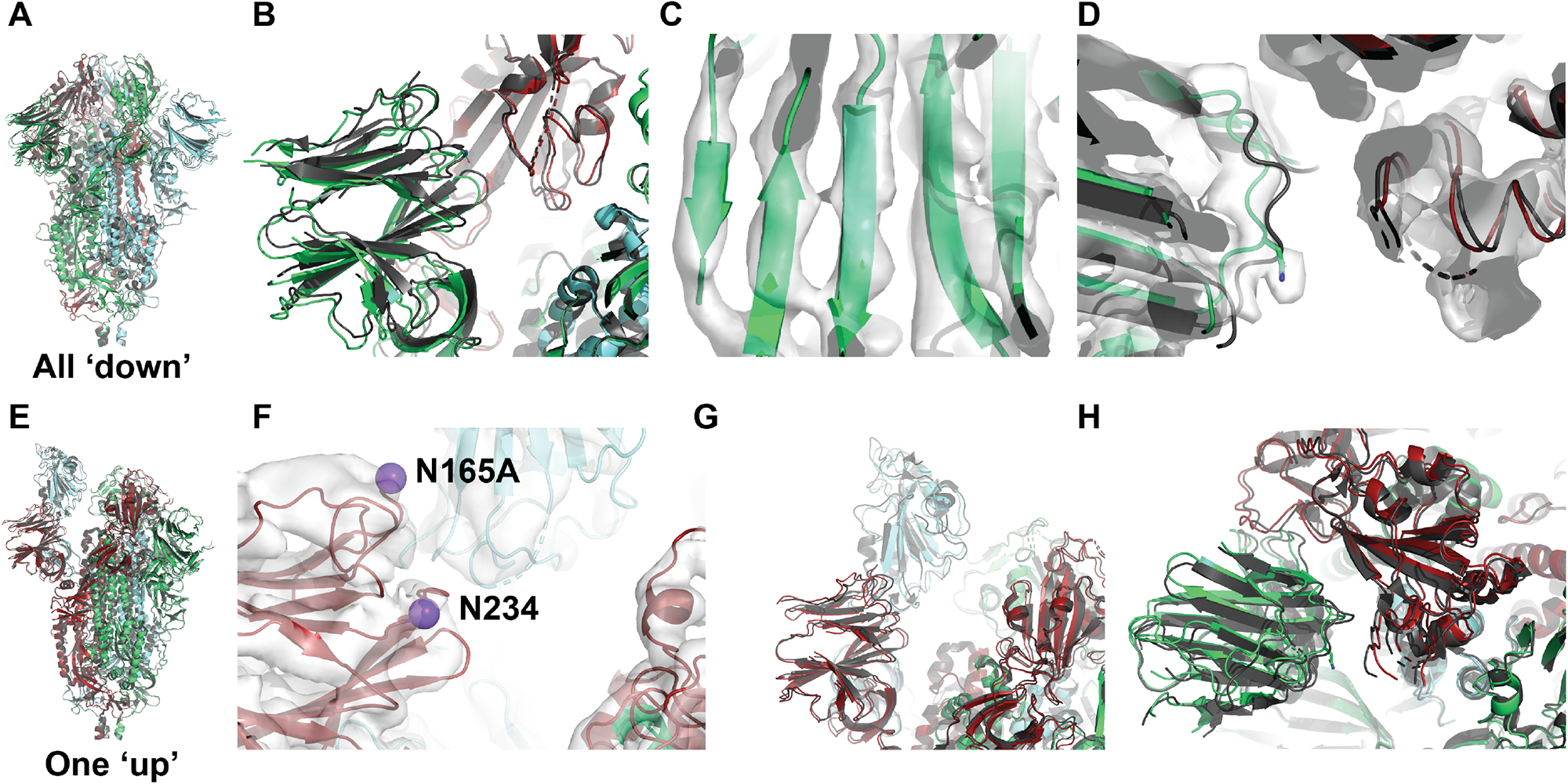
Structural comparison of the N165A mutant in the ‘up’ and ‘down’ configurations to the unmutated spike. **A)** Side view of the symmetric ‘down’ state N165A mutant S-protein trimer aligned to the unmutated trimer (PDB ID 6VXX, grey). **B)** Side view of a single NTD (green) and adjacent RBD (red) with the ‘down’ state structure (grey) depicting the shift in the position of the NTD. **C)** Map view of the apical β-sheet motif of the NTD for the N165A ‘down’ state **D)** Zoomed in view of the NTD as in (*C*) with the map identifying the NTD shift. **E)** Side view of the symmetric ‘up’ state N165A mutant S-protein trimer aligned to the unmutated trimer (PDB ID 6VYB, grey). **F)** Side view of the ‘up’ state adjacent NTD (red) with N165 and N234 alpha carbons represented as spheres. **G)** Zoomed out view of the NTD as in (F) depicting the alignment of unmutated spike (grey) with the RBD in cyan. **H)** View of the NTD adjacent to the ‘down’ free state RBD.

## Discussion

Viral fusion proteins are often heavily glycosylated with the SARS-2 S-protein being no exception. Though decorated with fewer glycans than the HIV-1 Envelope protein, with 22 glycans per protomer^16^, the SARS-2 spike is well shielded from immune surveillance. The SARS-2 spike protein has proven remarkably sensitive to domain-domain interfacial mutations^12–15,19^ which led us to ask whether glycans near the NTD-RBD interface might also impact the configuration of the spike. Here, we have investigated the role of two NTD glycans at positions 234 and 165 in modulating S protein conformational dynamics by tracking the shift of RBD disposition in glycan-deleted mutants using binding to ACE-2 receptor, NSEM and cryo-EM analysis. While the specific magnitudes of differences vary between the different analysis methods, the results tend to track in the same direction showing that deletion of glycan 234 shifts the RBD dynamics more toward the “down” state, whereas deletion of glycan 165 appears to slightly enhance the distribution toward more “up” states. The shift in the position of the NTD toward the RBD in the ‘down’ state N234A mutant suggest the N234 glycan plays a direct role in destabilizing the ‘down’ state RBD position such that removal allows tighter packing of the RBD to the NTD. Additionally, the observed shift in the position of the ‘up’ state RBD suggests a role for the N234 glycan in modulating RBD stability. This is consistent with a recently released theoretical study investigating ‘up’ state RBD sensitivity to the presence of N165/N234 glycans via molecular simulation^20^. This investigation found that the absence of these glycans resulted in a comparatively unstable ‘up’ state RBD. The results here confirm the prediction from these simulations that loss of the N234 glycan results in an increased prevalence of the ‘down’ state. Deletion of the glycan at position 165 here suggests an opposite effect on the conformation of the spike relative to the N234A mutant, with the NTD shifting away from the adjacent RBD. Though this appears to relieve strain caused by the restriction imposed by the N234 glycan, the resultant lack of packing between the RBD and NTD may be sufficient to favor transitions to the ‘up’ state. Further, this shift suggests the N165 glycan interacts directly with the RBD. Though direct interactions are not observed in the cryo-EM densities here or in previously published SARS-2 structures, the presence of ‘down’ state conformational heterogeneity evinced by the poor resolution of the RBD and NTD elements of the spike is consistent with the possibility of such an interaction. A more detailed examination of this heterogeneity and the influence of these glycans on the various states of the spike will likely require large datasets with improved orientational sampling to better resolve these apical regions. Nevertheless, the results here demonstrate that the conformational ensemble of the SARS-2 spike, and likely all β-CoV spikes, is sensitive to glycosylation patterns, especially near the NTD-RBD interface.

The results from this study lend insights into two key questions – what role may the glycans at positions 165 and 234 play in modulating RBD dynamics of the native SARS-2 spike and how might these findings impact vaccine design? Toward the first question, we recognize that the results we describe are in the context of a stabilized, ectodomain construct and differences between these and what occurs on the spike in its native context can be expected. Indeed, a recent report for a detergent solubilized, full-length SARS-2 spike suggested greater stability in the ‘down’ state RBD^21^. Yet our experimental results revealing the role of the N165 and N234 glycans in modulating the conformational landscape of the S protein, taken together with the findings from the computational analysis performed in the context of the full-length spike^20^, and our analysis of the RBD-proximal NTD glycans of diverse β-CoVs (Figure 1), provides strong support for a role for these glycans in controlling the virion bound S-protein conformation and dynamics. The differences in glycosylation in this region in different CoV spikes could be a contributor to determining their receptor specificity and thus their transmission. Toward the second question related to the utility for vaccine design, building upon our previous study where we demonstrated conformational control of RBD dynamics in the S protein ectodomain by modulating inter-domain protein-protein contacts, here we expand the tools for achieving such control to glycan-protein interactions, and demonstrate that RBD dynamics can be modulated by targeting key glycans at interdomain contacts. In so doing, we create two new ectodomain constructs with differential exposure of the immunodominant RBD for use as immunogens in vaccination regimens. Taken together, these investigations further demonstrate the remarkable plasticity of this conformational machine and suggest the S-protein has a diverse landscape of conformational escape mutations from which to select as genetic drift and host immune pressures direct its evolution.

**Table 1:** Cryo-EM data collection, refinement and validation statistics.

|  | N234A 'Down'<br>(EMD-XXXXX,<br>PDB YYYY) | N234A One<br>'Up'<br>(EMD-XXXXX,<br>PDB YYYY) | N165A 'Down'<br>(EMD-XXXXX,<br>PDB YYYY) | N165A One<br>'Up'<br>(EMD-XXXXX,<br>PDB YYYY) |
| --- | --- | --- | --- | --- |
| Data collection and processing |  |  |  |  |
| Microscope | Krios | Krios | Krios | Krios |
| Magnification | 81,000 | 81,000 | 75,000 | 75,000 |
| Voltage (kV) | 300 | 300 | 300 | 300 |
| Electron exposure (e-/Å<sup>2</sup>) | 64.97 | 64.97 | 47.51 | 47.51 |
| Detector | K3 | K3 | Falcon | Falcon |
| Defocus range (µm) | 0.44-4.63 | 0.44-4.63 | 0.39-2.70 | 0.39-2.70 |
| Pixel size (Å) | 1.065 | 1.065 | 1.050 | 1.050 |
| Micrographs Collected | 8,068 | 8,068 | 7,269 | 7,269 |
| Symmetry imposed | C3 | C1 | C3 | C1 |
| Initial particle images (no.) | 2,066,151 | 2,066,151 | 2,120,801 | 2,120,801 |
| Final particle images (no.) | 878,645 | 60,592 | 142,576 | 122,512 |
| Map resolution (Å) | 3.0 | 4.8 | 3.3 | 3.6 |
| FSC threshold | 0.143 | 0.143 | 0.143 | 0.143 |
| Map resolution range (Å) | 2.60-4.89 | 4.13-32.63 | 2.69-6.89 | 3.04-8.30 |
| Refinement |  |  |  |  |
| Initial model used | PDB 6VXX | PDB 6VYB | PDB 6VXX | PDB 6VYB |
| Map sharpening B factor (Å<sup>2</sup>) | -147.6 | -192.1 | -144.4 | -147.3 |
| R.m.s. deviations |  |  |  |  |
| Bond lengths (Å) | 0.015 | 0.012 | 0.012 | 0.011 |
| Bond angles (°) | 1.374 | 1.896 | 1.899 | 1.883 |
| Validation |  |  |  |  |
| MolProbity score | 1.63 | 1.58 | 1.05 | 1.07 |
| Clashscore | 4.44 | 0.64 | 0.27 | 0.24 |
| Poor rotamers (%) | 0.59 | 2.70 | 0.70 | 0.84 |
| Ramachandran plot |  |  |  |  |
| Favored (%) | 93.78 | 90.39 | 93.04 | 92.21 |
| Allowed (%) | 6.22 | 8.97 | 6.43 | 7.19 |
| Disallowed (%) | 0.00 | 0.64 | 0.53 | 0.61 |

## Acknowledgements

Cryo-EM data were collected on the Titan Krios at the Shared Materials and Instrumentation Facility in Duke University, and at the National Center for Cryo-EM Access and Training (NCCAT) and the Simons Electron Microscopy Center located at the New York Structural Biology Center, supported by the NIH Common Fund Transformative High Resolution Cryo-Electron Microscopy program (U24 GM129539,) and by grants from the Simons Foundation (SF349247) and NY State. We thank Ed Eng, Carolina Hernandez, Mark Walters and Holly Leddy for microscope alignments and assistance with cryo-EM data collection, and Alberto Bartesaghi for on-the-fly cryo-EM data pre-processing implemented on the Duke Titan Krios. This work was supported by UM1 AI100645 (B.F.H), the Duke Center for HIV/AIDS Vaccine Immunology-Immunogen Discovery, and UM1 AI44371 (B.F.H.), the Duke Consortium for HIV/AIDS Vaccine Development, Division of AIDS, NIAID, NIH; Duke University Center for AIDS Research (CFAR); R01AI145687 and Translating Duke Health Initiative (P.A. and R.C.H.). This study utilized the computational resources offered by Duke Research Computing (http://rc.duke.edu; NIH 1S10OD018164-01) at Duke University. We thank M. DeLong, C. Kneifel, M. Newton, V. Orlikowski, T. Milledge, and D. Lane from the Duke Office of Information Technology and Research Computing for providing assistance with setting up and maintaining the computing environment.

## Author contributions

R.C.H and P.A. conceived the designed the study, and analyzed data. R.C.H. processed cryo-EM data, and performed map and coordinate refinement. R.J.E. and K.M. performed NSEM studies. K.J. and V.S. produced and purified proteins. M.K. optimized cryo-EM specimen and collected cryo-EM data. R.J.E., M.B., B.F.H and P.A. supervised studies. R.C.H and P.A. wrote the manuscript with help from all authors.

## Data Availability

Cryo-EM reconstructions were deposited in the Electron Microscopy Data Bank with EMDB accession codes <TBD> and in Protein Data Bank with PDB IDs <TBD>.

## Methods

### Protein expression and purification

The SARS-CoV-2 ectodomain constructs were produced and purified as described previously^2^. Briefly, a gene encoding residues 1-1208 of the SARS-CoV-2 S (GenBank: MN908947) with proline substitutions at residues 986 and 987, a “GSAS” substitution at the furin cleavage site (residues 682–685), a C-terminal T4 fibritin trimerization motif, an HRV3C protease cleavage site, a TwinStrepTag and an 8XHisTag was synthesized and cloned into the mammalian expression vector pαH. All mutants were introduced in this background. Expression plasmids encoding the ectodomain sequence were used to transiently transfect FreeStyle293F cells using Turbo293 (SpeedBiosystems). Protein was purified on the sixth day posttransfection from the filtered supernatant using StrepTactin resin (IBA).

The ACE-2 gene was cloned as a fusion protein with a mouse Fc region attached to its C-terminal end. A 6X His-tag was added to the C-terminal end of the Fc domain. ACE-2 with mouse FC tag was purified by Ni-NTA chromatography.

### Thermal shift assay

The thermal shift assay was performed using Tycho NT. 6 (NanoTemper Technologies). Spike variants were diluted (0.15 mg/ml) in nCoV buffer (2mM Tris, pH 8.0, 200 mM NaCl, 0.02% sodium azide) and run in duplicates in capillary tubes. Intrinsic fluorescence was recorded at 330 nm and 350 nm while heating the sample from 35-95 °C at a rate of 3 °C/min. The ratio of fluorescence (350/330 nm) and the Ti were calculated by Tycho NT. 6.

### Cryo-EM sample preparation, data collection and processing

Purified SARS-CoV-2 spike preparations were diluted to a concentration of ~1 mg/mL in 2 mM Tris pH 8.0, 200 mM NaCl and 0.02% NaN3. 2.5 μL of protein was deposited on a CF-1.2/1.3 grid that had been glow discharged for 30 seconds in a PELCO easiGlow™ Glow Discharge Cleaning System. After a 30 s incubation in >95% humidity, excess protein was blotted away for 2.5 seconds before being plunge frozen into liquid ethane using a Leica EM GP2 plunge freezer (Leica Microsystems). Frozen grids for the N234A mutant were imaged in a Titan Krios (Thermo Fisher) equipped with a K3 detector (Gatan) while those for the N165A mutant were image in a Titan Krios equipped with a Falcon 3EC detector. Data were acquired using the Leginon system^23^ (N234A) or the EPU software (Thermo Fisher Scientific; N165A). The dose was fractionated over 50 raw frames and collected at 50ms framerate. This dataset was energy-filtered with a slit width of 30 eV for the N234A mutant. Individual frames were aligned and dose-weighted^24^. CTF estimation, particle picking, 2D classifications, *ab initio* model generation, heterogeneous refinements, homogeneous 3D refinements and local resolution calculations were carried out in cryoSPARC^25^.

### Cryo-EM structure fitting and analysis

Structures of the all ‘down’ state (PDB ID 6VXX) and single RBD ‘up’ state (PDB ID 6VYB) from the previously published SARS-CoV-2 ectodomain were used to fit the cryo-EM maps in Chimera^26^. Mutations were made in PyMol^27^. Coordinates were fit to the maps first using ISOLDE^28^ followed by iterative refinement using Phenix^29^ real space refinement and subsequent manual coordinate fitting in Coot as needed. Structure and map analysis were performed using PyMol, Chimera^26^ and ChimeraX^30^.

### Surface Plasmon Resonance

The binding of ACE-2 to the SARS-2 spike constructs was assessed by surface plasmon resonance on Biacore T-200 (GE-Healthcare) at 25°C with HBS-EP+ (10 mM HEPES, pH 7.4, 150 mM NaCl, 3 mM EDTA, and 0.05% surfactant P-20) as the running buffer. ACE-2 tagged at its C-terminal end to a mouse Fc region was captured on an anti-Fc surface. Binding was assessed by flowing over different concentrations of the spike constructs over the ACE-2 surface. The surface was regenerated between injections by flowing over 3M MgCl2 solution for 10s with flow rate of 100μl/min. Blank sensorgrams were obtained by injection of the same volume of HBS-EP+ buffer in place of IgGs and Fab solutions. Sensorgrams were corrected with corresponding blank curves. Sensorgram data were analyzed using the BiaEvaluation software (GE Healthcare).

