## Supplemental Data for "Glycans on the SARS-CoV-2 Spike Control the Receptor Binding Domain Conformation"

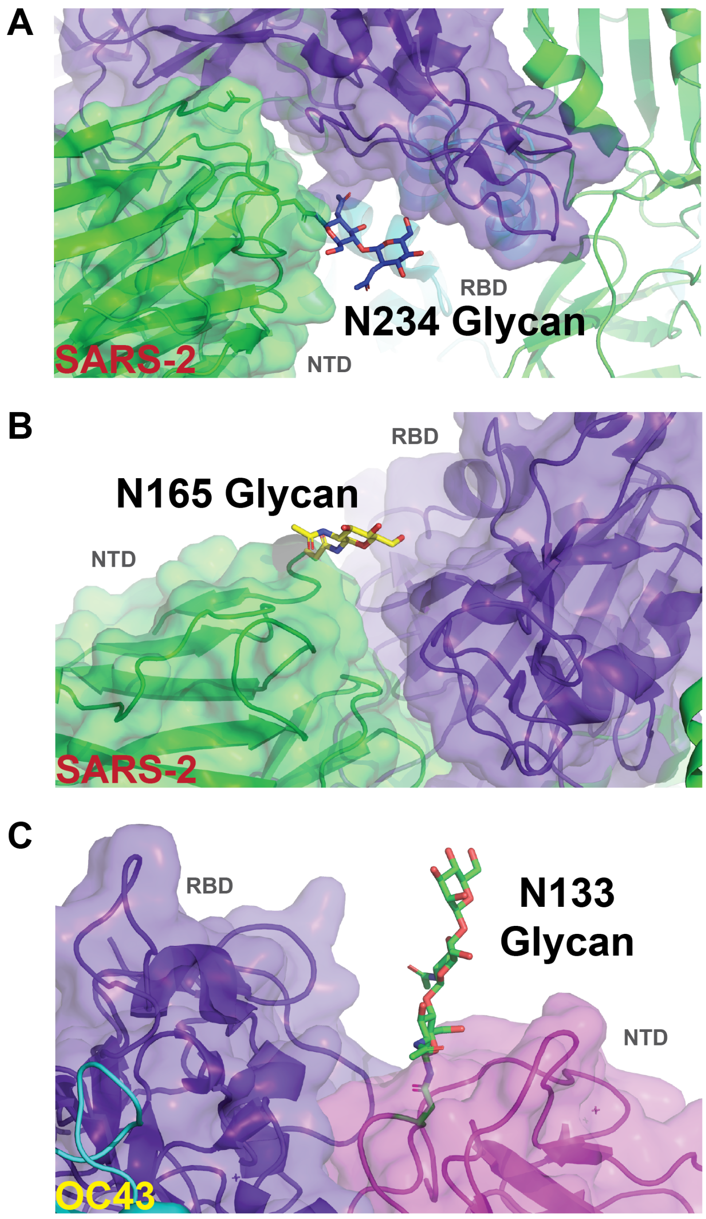


**Supplemental Figure 1. SARS-2 and OC43 RBD proximal NTD glycans. A)** Side view of a SARS-2 NTD (green) and RBD (purple) structure (PDB ID 6VXX) depicting the N234 glycan cleft. An RBD only structure (PDB ID 6M0J) was aligned to the trimer as a portion of the RBD is not present in the trimer structure. **B)** Side view of a SARS-2 NTD (green) and RBD (purple) structure (PDB ID 6VXX) depicting the N165 glycan. **C)** Side view of an OC43 NTD (magenta) and RBD (purple) structure (PDB ID 6OHW) depicting the N133 glycan.

A

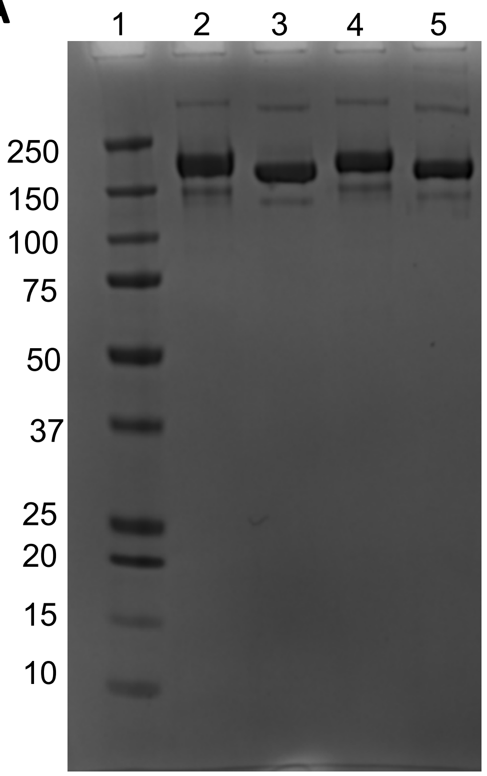


Legend:
1. Marker
2. Reduced N234A (93KJ)
3. Non-Reduced N234A (93KJ)
4. Reduced N165A (94KJ)
5. Non-Reduced N165A (94KJ)

| Plasmid | Yield (mg) |
|---|---|
| N234A | 2.0 |
| N165A | 0.8 |

B

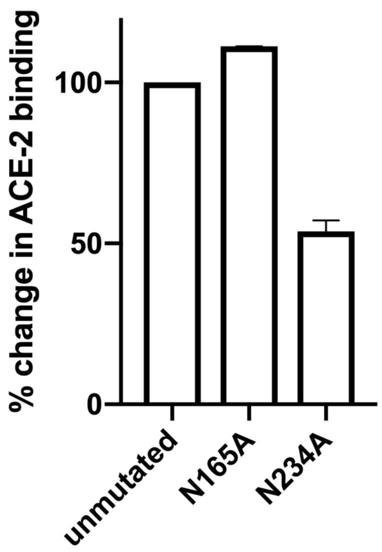


C

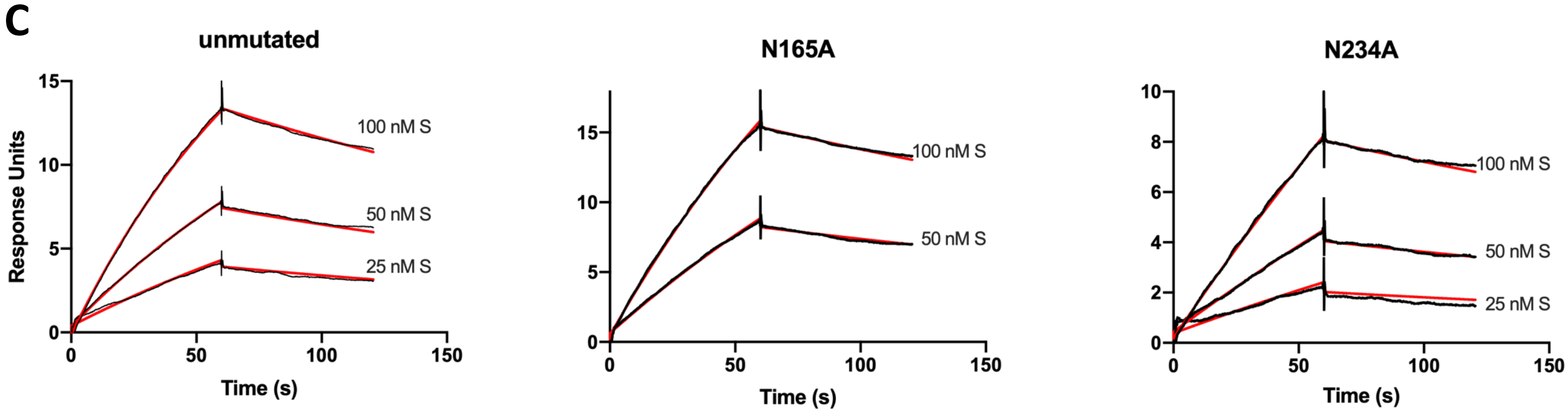


| | ka ($M^{-1}s^{-1}$) | Kd ($s^{-1}$) | KD (M) |
|---|---|---|---|
| Unmutated | 7.869E+4 | 0.003600 | 4.575E-8 |
| N165A | 4.792E+4 | 0.002730 | 5.697E-8 |
| N234A | 3731 | 0.002853 | 7.648E-7 |

**Supplemental Figure 2. SDS-PAGE and yields of purified S protein constructs. A)** SDS-PAGE gels of the S protein constructs. R= reducing conditions; NR = non-reducing conditions and expression yields/L of the S protein constructs. **B)** Independent SPR replicate measures for the unmutated, N164A, and N234A mutants. Error bars represent results from multiple injections. **C)** Binding of unmutated, N165A, and N234A S-protein to ACE-2. ACE-2 was captured on an anti-Fc surface via a C-terminal Fc tag, and binding was measured by flowing over different concentrations of the spike constructs in independent injections. Binding curves are shown in the upper panel. Black lines show the double-referenced binding data and the red lines show the global fit of the data to a Langmuir 1:1 binding model. Affinity and kinetics values are shown in the lower panel.

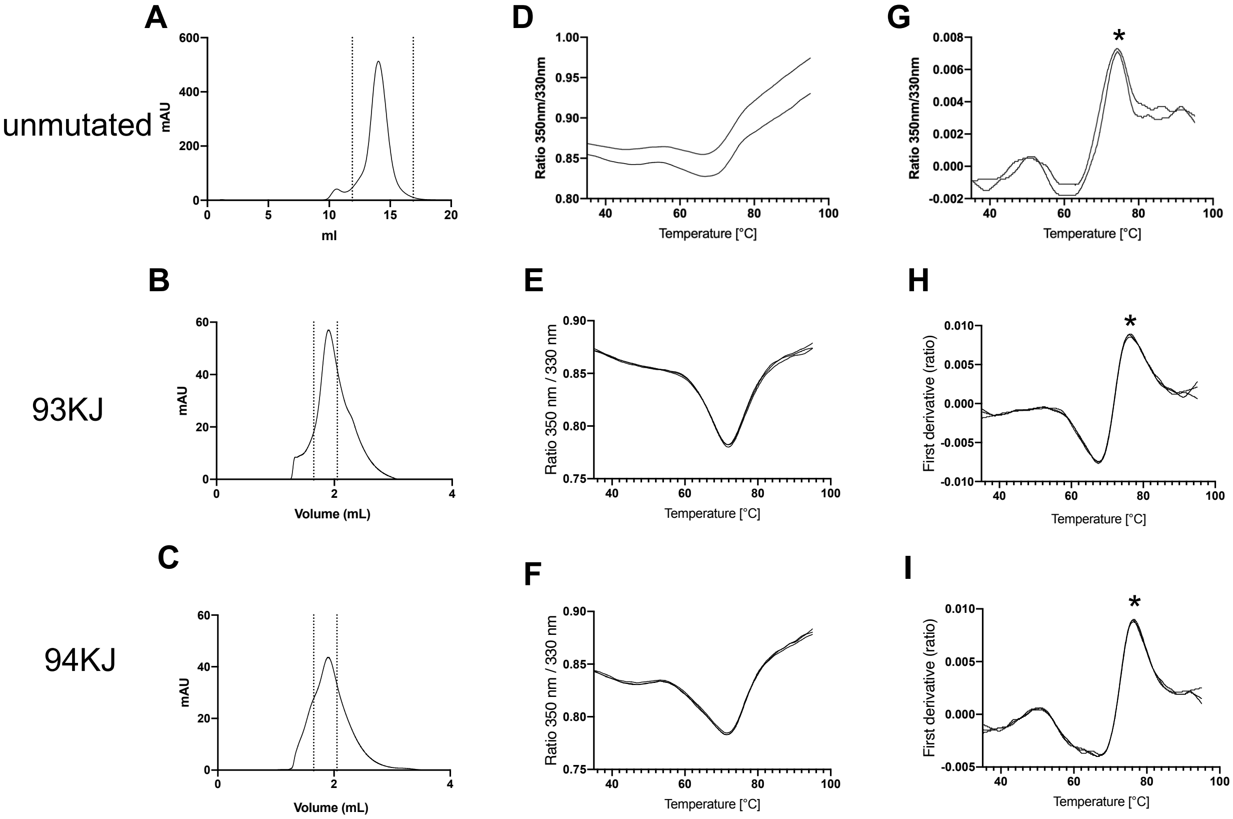


**J**

| | Ti value (℃) |
|---|---|
| Unmutated | 74.3 |
| N234A | 76.26 |
| N165A | 76.33 |

[a]Ti, inflection temperature determined at in the nCoV buffer (2 mM Tris pH 8.00, 200 mM NaCl, 0.02% sodium azide).

**Supplemental Figure 3. Thermostability of the S protein constructs. A-C)** SEC profile of the S proteins. The dotted lines indicate the portion of the peak that was collected for further studies. The unmutated spike was run on a Superose 6 Increase 10/300 column, and N234A and N165A spike was run on an analytical Superose 6 Increase 5/150 column. **D-I)** Unfolding profile curves obtained by intrinsic fluorescence measurements using Tycho NT. 6. **D-F)** show ratio between fluorescence at 350 nm and 330 nm. **G-I)** plot the first derivative of this ratio. Asterisk mark the inflection temperatures that are tabulated in **J)**

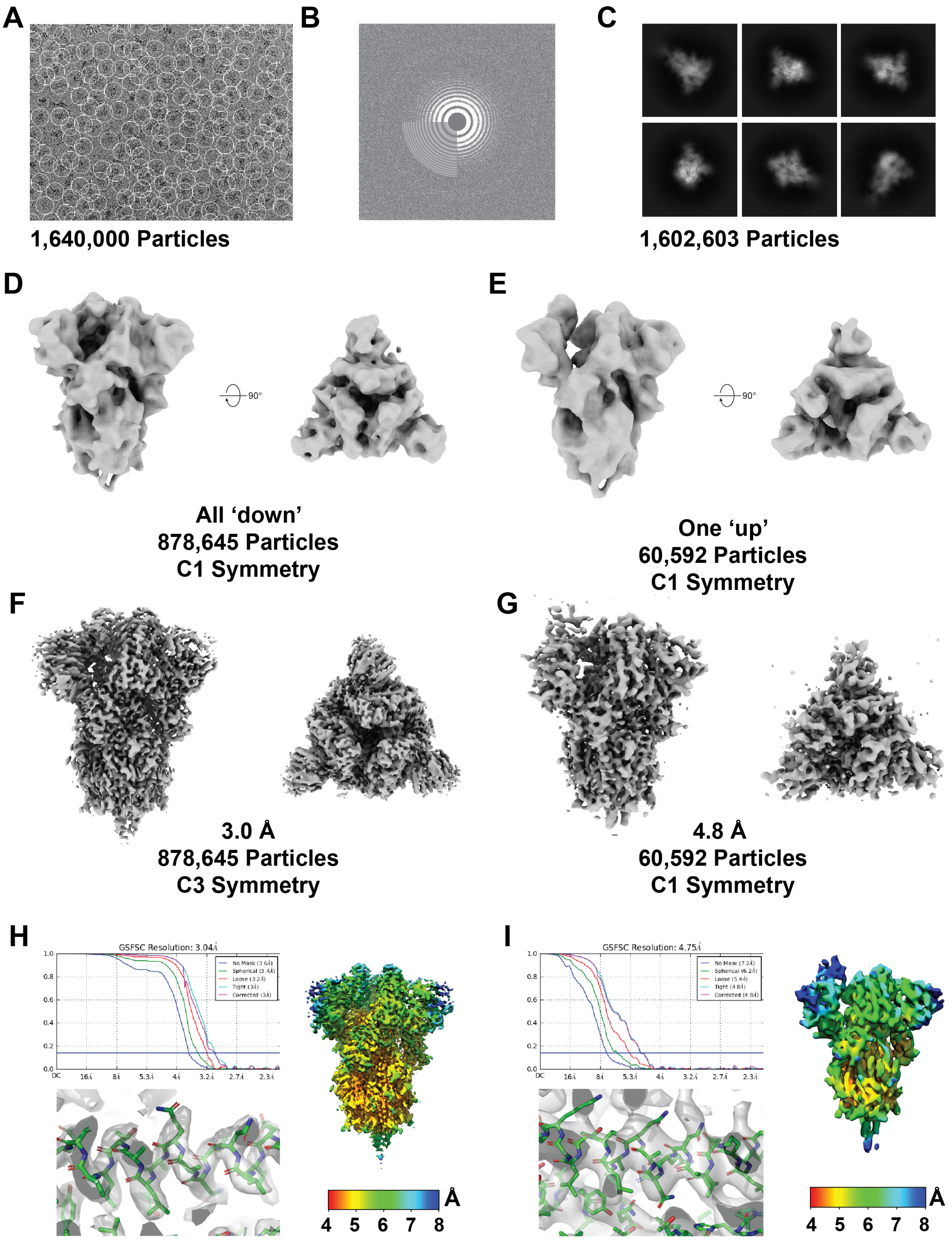

A
1,640,000 Particles
B
C
1,602,603 Particles
D
90°
All ‘down’
878,645 Particles
C1 Symmetry
E
90°
One ‘up’
60,592 Particles
C1 Symmetry
F
3.0 Å
878,645 Particles
C3 Symmetry
G
4.8 Å
60,592 Particles
C1 Symmetry
H
GSFSC Resolution: 3.04Å
No Mask (3.6Å)
Spherical (3.4Å)
Loose (3.2Å)
Tight (3Å)
Corrected (3Å)
1.0
0.8
0.6
0.4
0.2
0.0
DC
16Å
8Å
5.3Å
4Å
3.2Å
2.7Å
2.3Å
Å
4 5 6 7 8
I
GSFSC Resolution: 4.75Å
No Mask (7.2Å)
Spherical (6.2Å)
Loose (5.4Å)
Tight (4.8Å)
Corrected (4.8Å)
1.0
0.8
0.6
0.4
0.2
0.0
DC
16Å
8Å
5.3Å
4Å
3.2Å
2.7Å
2.3Å
Å
4 5 6 7 8

**Supplemental Figure 4. High-resolution cryo-EM structure determination pipeline for the N234A mutant ‘up’ and ‘down’ states. A)** Representative micrograph with selected particles circled. **B)** Representative CTF fit. **C)** Representative 2D classes **D)** Ab initio reconstruction of the ‘down’ state trimer depicting side (left) and top (right) views. **E)** Ab initio reconstruction of the ‘up’ state trimer depicting side (left) and top (right) views. **F)** High-resolution map of the C3 symmetric refinement of the ‘down’ state depicting side (left) and top (right) views. **G)** High-resolution map of the C1 asymmetric refinement of the ‘up’ state depicting side (left) and top (right) views. **H)** (*top left*) Fourier shell correlation curve for the ‘down’ state map, (*bottom left*) representative density, (*right*) local map resolutions. **I)** (*top left*) Fourier shell correlation curve for the ‘up’ state map, (*bottom left*) representative density, (*right*) local map resolutions.

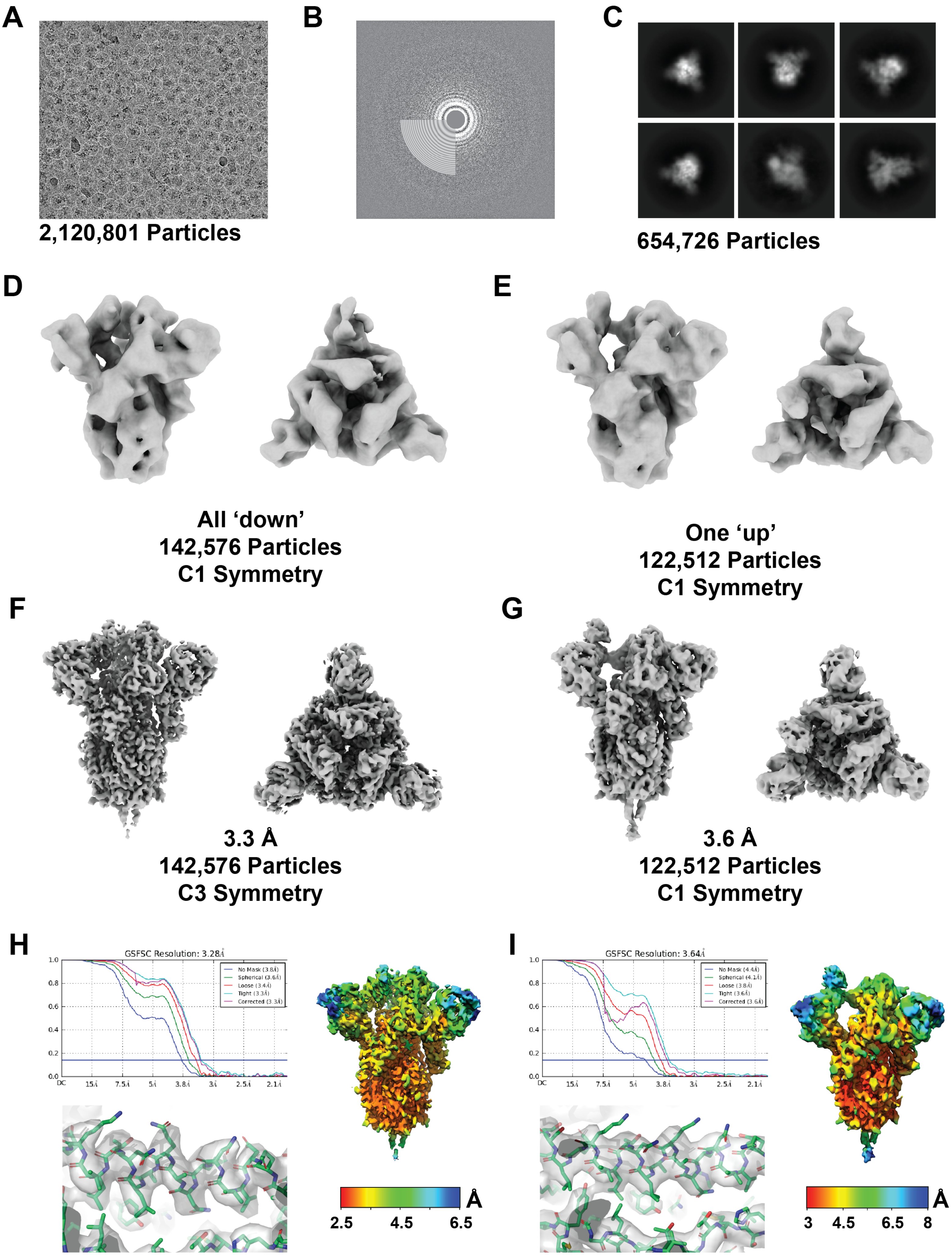
A
2,120,801 Particles
B
C
654,726 Particles
D
All ‘down’
142,576 Particles
C1 Symmetry
E
One ‘up’
122,512 Particles
C1 Symmetry
F
3.3 Å
142,576 Particles
C3 Symmetry
G
3.6 Å
122,512 Particles
C1 Symmetry
H
GSFSC Resolution: 3.28Å
No Mask (3.8Å)
Spherical (3.6Å)
Loose (3.4Å)
Tight (3.3Å)
Corrected (3.3Å)
1.0
0.8
0.6
0.4
0.2
0.0
DC
15Å
7.5Å
5Å
3.8Å
3Å
2.5Å
2.1Å
Å
2.5
4.5
6.5
I
GSFSC Resolution: 3.64Å
No Mask (4.4Å)
Spherical (4.1Å)
Loose (3.8Å)
Tight (3.6Å)
Corrected (3.6Å)
1.0
0.8
0.6
0.4
0.2
0.0
DC
15Å
7.5Å
5Å
3.8Å
3Å
2.5Å
2.1Å
Å
3
4.5
6.5
8

**Supplemental Figure 5. High-resolution cryo-EM structure determination pipeline for the N165A mutant 'up' and 'down' states. A)** Representative micrograph with selected particles circled. **B)** Representative CTF fit. **C)** Representative 2D classes **D)** Ab initio reconstruction of the 'down' state trimer depicting side (left) and top (right) views. **E)** Ab initio reconstruction of the 'up' state trimer depicting side (left) and top (right) views. **F)** High-resolution map of the C3 symmetric refinement of the 'down' state depicting side (left) and top (right) views. **G)** High-resolution map of the C1 asymmetric refinement of the 'up' state depicting side (left) and top (right) views. **H)** (*top left*) Fourier shell correlation curve for the 'down' state map, (*bottom left*) representative density, (*right*) local map resolutions. **I)** (*top left*) Fourier shell correlation curve for the 'up' state map, (*bottom left*) representative density, (*right*) local map resolutions.

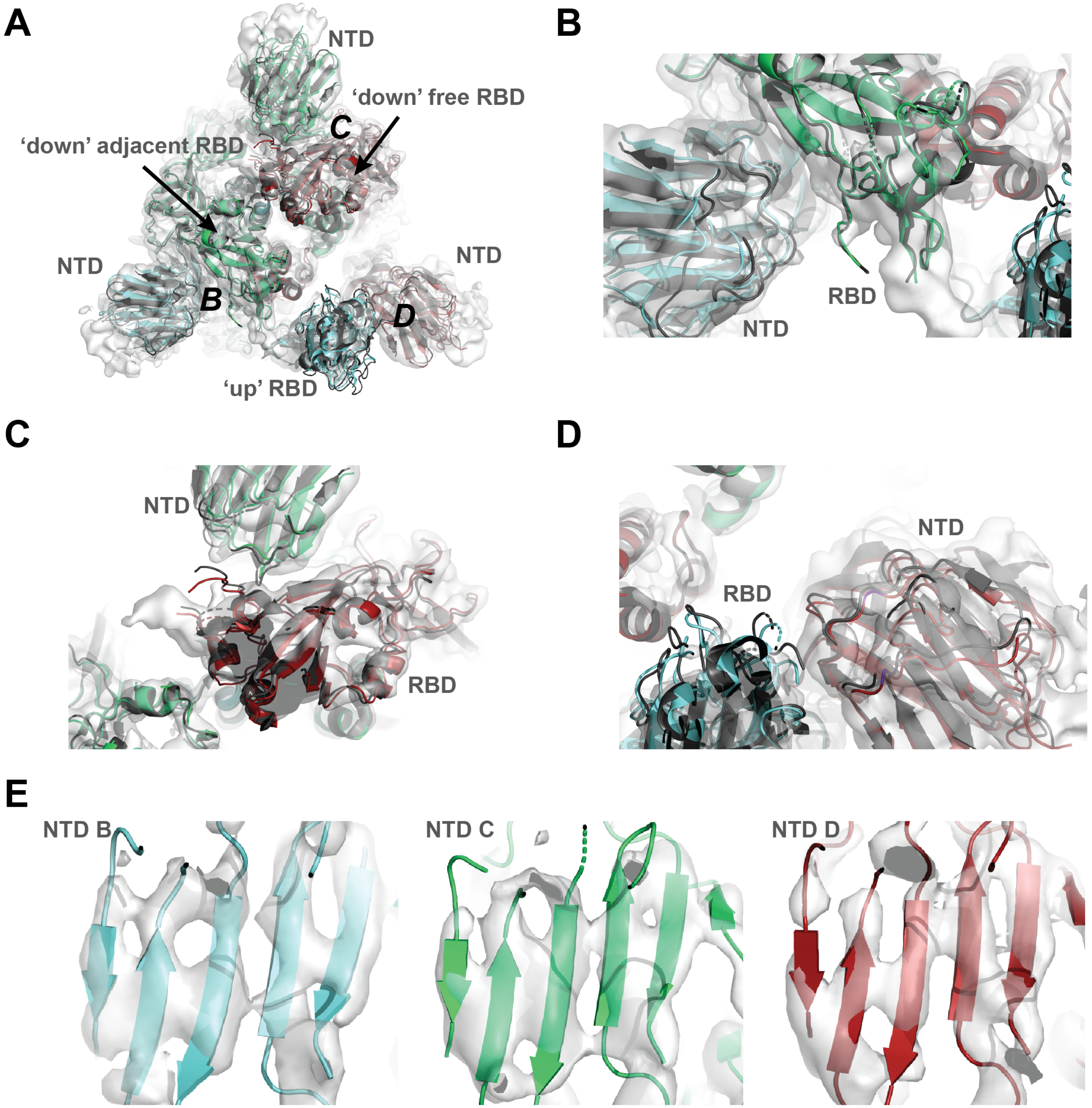


**Supplemental Figure 6. Structure of the 'up' state N165A mutant NTD shifts. A)** Top view of the 'up' state N165A structure (green, cyan, and red) aligned to the unmuated spike (PDB ID 6VYB, grey). **B)** View of the 'down' adjacent RBD (green) and NTD (cyan) aligned to the unmutated spike (grey). **C)** View of the 'down' free RBD (red) and NTD (green) aligned to the unmutated spike (grey). **D)** View of the 'up' state RBD (cyan) and NTD (red) aligned to the unmutated spike (grey). **E)** Map view of the apical β-sheet motifs of the NTDs for the N165A 'up' state
